# Metabolomic analysis of *Drosophila melanogaster* larvae lacking Pyruvate kinase

**DOI:** 10.1101/2023.06.05.543743

**Authors:** Yasaman Heidarian, Jason P. Tourigny, Tess D. Fasteen, Nader H. Mahmoudzadeh, Alexander J. Hurlburt, Travis Nemkov, Julie A. Reisz, Angelo D’Alessandro, Jason M. Tennessen

## Abstract

Pyruvate kinase (Pyk) is a rate-limiting enzyme that catalyzes the final metabolic reaction in glycolysis. The importance of this enzyme, however, extends far beyond ATP production, as Pyk is also known to regulate tissue growth, cell proliferation, and development. Studies of this enzyme in *Drosophila melanogaster*, however, are complicated by the fact that the fly genome encodes six Pyk paralogs whose functions remain poorly defined. To address this issue, we used sequence distance and phylogenetic approaches to demonstrate that the gene *Pyk* encodes the enzyme most similar to the mammalian Pyk orthologs, while the other five *Drosophila* Pyk paralogs have significantly diverged from the canonical enzyme. Consistent with this observation, metabolomic studies of two different *Pyk* mutant backgrounds revealed that larvae lacking Pyk exhibit a severe block in glycolysis, with a buildup of glycolytic intermediates upstream of pyruvate. However, our analysis also unexpectedly reveals that steady state pyruvate levels are unchanged in *Pyk* mutants, indicating that larval metabolism maintains pyruvate pool size despite severe metabolic limitations. Consistent with our metabolomic findings, a complementary RNA-seq analysis revealed that genes involved in lipid metabolism and peptidase activity are elevated in *Pyk* mutants, again indicating that loss of this glycolytic enzyme induces compensatory changes in other aspects of metabolism. Overall, our study provides both insight into how Drosophila larval metabolism adapts to disruption of glycolytic metabolism as well as immediate clinical relevance, considering that Pyk deficiency is the most common congenital enzymatic defect in humans.

## INTRODUCTION

The enzyme Pyruvate kinase (Pyk; E.C. 2.7.1.40) catalyzes the final reaction of glycolysis (phosphoenolpyruvate + ADP + Pi to pyruvate + ATP). Since the Pyk-catalyzed reaction is essentially irreversible under normal cellular conditions (SCHORMANN *et al*. 2019), this enzyme controls a key rate-limiting, “pay-off” glycolytic step that controls ATP production, and dictates the rate at which glycolytic intermediates enter the metabolic pathways that branch off both glycolysis and the TCA cycle. In so doing, Pyk regulates fluxes towards serine synthesis, which indirectly influences redox balance and nucleoside anabolism via the amino acids cysteine (de novo glutathione synthesis) and glycine (one-carbon metabolism for de novo purine synthesis) (MADDOCKS *et al*. 2013). Thus, changes in Pyk activity can influence cell growth, proliferation, and differentiation by balancing the flux of glycolytic metabolites between mitochondrial energy production and the synthesis of metabolites involved in chromatin remodeling, one-carbon metabolism, and biomass production (ISRAELSEN AND VANDER HEIDEN 2015; HSU AND HUNG 2018).

The importance of Pyk in human health is clear from the myriad of diseases that are caused by altered activity of the two human orthologs, Pyruvate Kinase, Liver and Red Blood Cell (PKLR) and Pyruvate Kinase, Muscle (PKM). For example, mutations in the *PKLR* gene result in a common congenital disorder known as Pyk deficiency (PKD) (FATTIZZO *et al*. 2022; LUKE *et al*. 2023). Individuals with PKD experience a wide spectrum of clinical and biochemical symptoms and have a highly variable prognoses that depend on environmental factors, early interventions, and the severity of the causal mutations (DOLAN *et al*. 2002; GRACE *et al*. 2015). Similarly, changes in splicing of PKM transcripts are well-documented in a number of disease states, with a switch from the PKM1 isoform towards the PKM2 isoform associated with pulmonary hypertension (CARUSO *et al*. 2017; ZHANG *et al*. 2017), sporadic Alzheimer’s disease (TRAXLER *et al*. 2022), and enhancement of the Warburg effect in cancer cells (ISRAELSEN AND VANDER HEIDEN 2015). Finally, population genetic studies support a model in which *PKLR* mutations are protective against malaria (MIN-OO et al. 2007; AYI et al. 2008; MACHADO et al. 2012; VAN BRUGGEN et al. 2015), suggesting that partial inhibition of the enzyme could be exploited toward anti-malarial efforts. Basic studies of PYK function in the context of endogenous metabolic networks thus hold the potential to inform better interventions for a wide range of human diseases.

The fruit fly *Drosophila melanogaster* is a powerful model for studying the function of metabolic enzymes within the context of both normal physiology and human disease models (DRUMMOND-BARBOSA AND TENNESSEN 2020; KIM *et al*. 2021). In this regard, previous studies of *Drosophila* Pyk have demonstrated roles for this enzyme in fly development and behavior, with reduced *Pyk* levels being associated with defects in muscle and wing development, neuron and glial function, and olfactory memory (TIXIER *et al*. 2013; VOLKENHOFF *et al*. 2015; WU *et al*. 2018; SPANNL *et al*. 2020). However, the effects of decreased Pyk activity in the fly have not been examined at a metabolic level. Moreover, the fly genome encodes six predicted Pyk paralogs (Pyk, CG7069, CG2964, CG7362, CG11249, and CG12229,GARAPATI *et al*. 2019; GRAMATES *et al*. 2022b), thus highlighting a need to characterize the *in vivo* functional potential of individual enzymes within this gene class.

Here, we use sequence identities and phylogenetics in combination with metabolomics and transcriptomics to analyze *Pyk* (FBgn0267385), which encodes the most abundantly and widely expressed member of the *Drosophila melanogaster* Pyk family (GRAVELEY *et al*. 2011). Our studies indicate that the gene *Pyk* encodes the protein most analogous to the mammalian PKM and PKLR enzymes. Subsequent metabolomic analysis of two genetically distinct *Pyk* mutant backgrounds reveals a severe block in glycolysis, with a buildup of glycolytic intermediates and a depletion of lactate, 2-hydroxyglutarate (2HG), and the amino acid proline. Moreover, the loss of Pyk activity also induced a significant upregulation of genes involved in lipid metabolism and protease activity. Overall, our study validates the presumed function of Pyk within *Drosophila* metabolism and identifies the transcriptional and metabolic networks that respond to loss of Pyk activity during larval development.

## METHODS

### *Drosophila* husbandry and larval collection

Fly stocks were maintained on Bloomington *Drosophila* Stock Center media at 25°C. Larvae were collected on molasses agar plates as previously described (LI AND TENNESSEN 2017; LI AND TENNESSEN 2018). Briefly, 50 virgin females and 25 males of the appropriate genotypes were placed in six-ounce plastic bottles (Genesee Scientific; Cat #: 32-130) with holes punched in the side using a 20-gauge needle. A 35 mm molasses agar plate (Corning 353001) with a smear of yeast paste on the surface was taped in place in the mouth of the bottle and the bottle inverted and placed in a 25°C incubator. The molasses agar/yeast plate was replaced at least once per day. For sample collection, eggs were collected on the molasses agar/yeast plate for 6 hours, individual plates were placed inside a 60 mm cell culture dish, and the collected embryos were placed in a 25°C incubator and allowed to develop for 48 hours. Larvae were collected at the early L2 stage for subsequent metabolomics analysis.

### Generation of *Pyk* alleles using CRISPR/Cas9

The *Pyk* deletions *Pyk^23^* and *Pyk^31^* were generated using a previously described approach for CRISPR/Cas9 mutagenesis (GRATZ *et al*. 2013; SEBO *et al*. 2014). Briefly, oligos containing gRNA sequences #1 (5’-GTGCCCCATGTGCGTCTGTC-3’) and #2 (5’-GCTGCTGGAGGCAGGTCCGA-3’) were inserted into pU6-BbsI-chiRNA (DGRC Stock #1362) and the resulting plasmids were independently injected into BDSC Stock #52669 (*y^1^ M{RFP[3xP3.PB] GFP[E.3xP3]=vas-Cas9.S}ZH-2A w^1118^*) by Rainbow Transgenics (Camarillo, CA). Injected females and F1 progeny were crossed to *w*; ry^506^ Dr^1^/TM3, P{Dfd-GMR-nvYFP}3, Sb^1^*. Putative F2 *Pyk*^Δ^*/ TM3, P{Dfd-GMR-nvYFP}3, Sb^1^* siblings were mated and the F3 generation screened for animals lacking the *TM3* balancer, indicating the presence of a lethal mutation. Any strain that failed to generate homozygous mutant adults was crossed with *Pyk^61^* mutants and the F1 progeny analyzed for failure to complement. Mutant strains that failed to complement *Pyk^61^* were further analyzed for the presence of *Pyk* mutations. *Pyk^23^* was isolated from injections using gRNA#1 and *Pyk^31^*was isolated from injections using gRNA#2. The deletions were identified by amplifying and sequencing a region of the *Pyk* gene using oligos 5’-CACGCACTTTGTTTACATCAGC-3’ and 5’-GCACCAGTCCACGGTAGAGA-3’.

### Generation of *Pyk* alleles using p-element excision

Both the *Pyk^60^ and Pyk^61^* alleles were generated using standard techniques for imprecise p-element excision (ROBERTSON *et al*. 1988; BELLEN *et al*. 2004). Briefly, virgin females containing the transposon insertions *Pyk^DG05605^/TM3, Sb^1^, Ser^1^* and *Pyk^EY10213^/TM3, Sb^1^, Ser^1^*(BDSC stock 20088) were independently crossed with *ry506 p{*Δ*2-3}99B* males. F1 virgin females lacking the TM3 balancer were crossed with *w*; ry^506^ Dr^1^/TM3, P{Dfd-GMR-nvYFP}3, Sb^1^* males (BDSC stock 23231). F2 males with white eyes were then crossed to *w*; ry^506^ Dr^1^/TM3, P{Dfd-GMR-nvYFP}3, Sb^1^* females and F3 *Pyk*^Δ^*/ TM3, P{Dfd-GMR-nvYFP}3, Sb^1^*siblings were used to establish mutant lines. The *Pyk^60^* allele was generated by the imprecise excision of *Pyk^EY21058^* and the *Pyk^61^* deletion was generated by the imprecise excision of *Pyk^DG05605^*. Endpoints of the deletions were mapped using a PCR-based approach. The *Pyk^61^*deletion was sequenced by isolating homozygous mutant larvae, amplifying across the deletion using the oligos 5’-GATTTCCTTCAGAGCATTTGCGTC-3’ and 5’-TTCACCGTGCAGCAAGACATC-3’ and sequencing the resulting PCR product. The *Pyk^61^* deletion is 6400 bp long starting with the sequence 5’-GATGCCTTTGTTGCCGCCCT-3’ and ending with the sequence 5’-GTATGAACTTCTCTCACGGC-3’ and includes an 18 bp insertion of the sequence 5’-AAGTTCAAGTTCTGGATT-3’. The *Pyk^60^* allele represents a large deletion or rearrangement between the sequences 5’-CTTCTGTTCTATCCGATTGCCGG-3’ and 5’-TTGACGCGCTCATTGGGTTTC-3’. However, we were unable to produce a PCR product spanning the lesion. The *Pyk^prec^* allele is a precise excision of *Pyk^DG05605^*. For all analyses, the *Pyk^60^/TM3, P{Dfd-GMR-nvYFP}3, Sb^1^ and Pyk^61^/TM3, P{Dfd-GMR-nvYFP}3, Sb^1^* strains were crossed together and trans-heterozygous *Pyk^60/61^* mutant larvae were selected for based on the absence of YFP.

Since the transposon excision events that produced the *Pyk^60^* and *Pyk^61^* deletion alleles also disrupted neighboring genes, we further analyzed the effects of these deletions on expression of neighboring genes. When these deletions are placed in *trans*, the resulting heterozygote lacks sequence corresponding to nearly the entire *Pyk* coding region and the 5’ exons of the neighboring *Pyk* homolog *CG7069*. Northern blot analysis of trans-heterozygous L2 larvae revealed that *Pyk^60/61^* larvae fail to accumulate detectable *Pyk* mRNA transcripts (Figure S1). The mRNA of neighboring genes *Polr3F* and *CG18596*, however, were present at similar levels in both *Pyk^Prec^* and *Pyk^60/61^*, indicating that the phenotypes arising from the *Pyk* mutant background are specific to loss of *Pyk* expression. Of note, the second pyruvate kinase homolog disrupted by these deletions, *CG7069*, is not expressed during the embryonic and larval stages, and since *Pyk^60/61^* larvae die prior to the L3 stage, the effects of these deletions on *CG7069* expression could not be examined.

### Northern Blot Analysis

RNA was extracted from L2 larvae with TriPure isolation reagent and northern blot analysis was conducted as previously described (KARIM AND THUMMEL 1991; SULLIVAN AND THUMMEL 2003). The following PCR oligos were used to generate northern blot probes: *Pyk*: 5’-GCTGACCACCAACAAGGAAT-3’ and 5’-GCACCAGTCCACGGTAGAGA-3’ *Polr3F*: 5’-ACCAACGATGACCTGACCAAG-3’ and 5’-ATTGTTTCCAGATCGGCCTCC-3’ *rp49*: 5’-ACAAATGTGTATTCCGACCACG-3’ and 5’-TCAAGATGACCATCCGCCCAG-3’

### Phylogenetic Analysis of Pyk orthologs and paralogs

Protein sequences for each of the Pyk homologs were retrieved from FlyBase (GRAMATES *et al*. 2022a), Saccharomyces Genome Database (ENGEL *et al*. 2014; WONG *et al*. 2023), WormBase (DAVIS *et al*. 2022), NCBI RefSeq (O’LEARY *et al*. 2016), TAIR (BERARDINI *et al*. 2015), and UniProt (2023). The longest verified isoform of each was selected for multiple sequence alignment (MSA; see Table S1 for protein sequence information), except for mosquito which was a predicted entry (PETCHAMPAI *et al*. 2019). For the sequence identity and similarity heatmaps, an ensemble MSA was performed via the M-Coffee web server (WALLACE *et al*. 2006; MORETTI *et al*. 2007; NGUYEN *et al*. 2015; MINH *et al*. 2020), running the default command: ‘t_coffee –in=Pyk_homologs_E_coli_out.fa Mpcma_msa Mmafft_msa Mclustalw_msa Mdialigntx_msa Mpoa_msa Mmuscle_msa Mprobcons_msa Mt_coffee_msa – output=score_html clustalw_aln fasta_aln score_ascii phylip –tree –maxnseq=150 – maxlen=2500 –case=upper –seqnos=off –outorder=input –run_name=result – multi_core=4 –quiet=stdout’. The resulting MSA was analyzed using the ‘seqidentity()’ function of the Bio3D R package to extract pairwise matrices (GRANT *et al*. 2006), which were visualized with corrplot (WEI AND V. 2021).

For the phylogenetic analysis, we first performed MSA with MAFFT v7.520 using the E-INS-I algorithm to tolerate potentially large gaps between conserved segments (KATOH AND STANDLEY 2013), with the command ‘mafft –maxiterate 1000 –thread 4 – genafpair Pyk_homologs_incl_Aa_and_E_coli_outgroup.fa > mafft_einsi_output.msa.fa’. We then reconstructed the phylogenetic tree with IQ-TREE 2 using the WAG model of protein substitution (WHELAN AND GOLDMAN 2001; NGUYEN *et al*. 2015; HOANG *et al*. 2018; MINH *et al*. 2020), run with the command ‘iqtree2 –s mafft_einsi_output.msa.fa –redo –B 1000 –T 4 –m WAG –prefix mafft_einsi_WAG’. The resulting phylogeny was rooted to *E. coli*, formatted, and visualized using FigTree v1.4.4 (RAMBAUT 2018).

### Triglyceride Assays and Nile Red Staining

Triglyceride levels in *Pyk* control and mutant L2 larvae were measured as previously described (TENNESSEN *et al*. 2014). Briefly, synchronized populations of embryos were collected for 4 hrs on a molasses agar plate with yeast paste smeared on the surface, as described (LI AND TENNESSEN 2017). Embryos were allowed to hatch on the surface of the collection plate and larvae were reared on the same plate in a 25 °C incubator. Sixty hours after egg-laying, heterozygous controls and homozygous mutant larvae were identified based on lack of GFP expression from the *TM3, P{Dfd-GMR-nvYFP}3, Sb^1^* balancer chromosome. Twenty-five larvae of the appropriate genotypes were then collected in 1.5 mL microfuge tubes, washed three times with phosphate-buffered saline pH 7.0 (PBS), homogenized in 100 µl of cold PBS + 0.05% Tween 20 (PBST), and heat-treated for 10 min at 70°C. The resulting homogenate was assayed for triglyceride and soluble protein levels as previously described (TENNESSEN *et al*. 2014).

Nile red staining was conducted on L2 fat bodies as previously described (GRÖNKE *et al*. 2005). Briefly, dissected tissues were fixed with 4% paraformaldehyde in PBS for 30 min and washed with PBST three times for 5 minutes each, and then incubated for 1 hr at room temperature in a 1:1000 dilution of Nile Red stock (10mg/mL in acetone) in PBS. Tissues were washed three times for 5 minutes each in PBST, mounted in Vectashield (Vector Laboratories), and imaged on a Leica SP8 Confocal at 568nm.

### Gas Chromatography Mass Spectrometry (GC-MS)-based Metabolomics

*Pyk* mutant and control larvae were collected at the early L2 stage (∼48 hours after egg-laying at 25°C) as previously described (LI AND TENNESSEN 2018). All samples collected for analysis represent biological replicates obtained from independent mating bottles and each sample contained 25 larvae. Analysis of the *Pyk^60^ and Pyk^61^* alleles was conducted by the University of Utah metabolomics core facility as previously described (COX *et al*. 2017). *Pyk^23^*^/31^ mutants and *Pyk^23/+^* control samples were analyzed by the University of Colorado Metabolomics core facility. Data were normalized to sample mass and internal standards.

### Ultra High-pressure Liquid Chromatography – Mass Spectrometry (UHPLC-MS)-based Metabolomics

UHPLC-MS metabolomics analyses were performed at the University of Colorado Anschutz Medical Campus, as previously described (NEMKOV *et al*. 2019). Briefly, the analytical platform employs a Vanquish UHPLC system (Thermo Fisher Scientific, San Jose, CA, USA) coupled online to a Q Exactive mass spectrometer (Thermo Fisher Scientific, San Jose, CA, USA). The (semi)polar extracts were resolved over a Kinetex C18 column, 2.1 x 150 mm, 1.7 µm particle size (Phenomenex, Torrance, CA, USA) equipped with a guard column (SecurityGuard^TM^ Ultracartridge – UHPLC C18 for 2.1 mm ID Columns – AJO-8782 – Phenomenex, Torrance, CA, USA) using an aqueous phase (A) of water and 0.1% formic acid and a mobile phase (B) of acetonitrile and 0.1% formic acid for positive ion polarity mode, and an aqueous phase (A) of water:acetonitrile (95:5) with 1 mM ammonium acetate and a mobile phase (B) of acetonitrile:water (95:5) with 1 mM ammonium acetate for negative ion polarity mode. The Q Exactive mass spectrometer (Thermo Fisher Scientific, San Jose, CA, USA) was operated independently in positive or negative ion mode, scanning in Full MS mode (2 μscans) from 60 to 900 m/z at 70,000 resolution, with 4 kV spray voltage, 45 sheath gas, 15 auxiliary gas. Calibration was performed prior to analysis using the Pierce^TM^ Positive and Negative Ion Calibration Solutions (Thermo Fisher Scientific).

### Statistical Analysis of Metabolomics Data

Both metabolomics datasets were analyzed using Metaboanalyst 5.0 (PANG *et al*. 2021), with data first preprocessed using log normalization and Pareto scaling.

### RNA-seq Analysis

RNA from *Pyk^23/+^* heterozygous controls and *Pyk^23/31^* mutants was extracted from L2 larvae with TriPure isolation reagent and further purified using a Qiagen RNeasy kit (Catalog #74004). RNA-seq was performed on three whole-fly biological replicates for *Pyk^23/+^* heterozygous controls and *Pyk^23/31^* mutants. Samples were paired-end sequenced on the NextSeq 550 platform 75 cycles to a depth of 15-20 million reads each, using standard Illumina TruSeq Stranded mRNA libraries, at the IU Center for Genomics and Bioinformatics.

RNA-seq read quality was assessed with FastQC (ANDREWS 2010) and MultiQC (EWELS *et al*. 2016), with adequate reads and no significant issues noted. Reads were pseudo-aligned and quantified using Kallisto v0.46.0 (BRAY *et al*. 2016), the *D. melanogaster* BDGP6.32 reference assembly and annotation retrieved through Ensembl (CUNNINGHAM *et al*. 2022), and the gffread utility from the Cufflinks suite to generate the transcriptome from the assembly and annotation (TRAPNELL *et al*. 2010).

Differential expression analysis was performed with DESeq2 v1.30.1 (LOVE *et al*. 2014) running in RStudio v1.4.1717 on R v4.0.4. Genes with average expression below 2 counts per sample (i.e. 48 counts per row) were filtered out before determining genes with significant expression difference in at least one sample with the likelihood-ratio test (LRT) and LRT *p*-value ≤ 0.05. Subsequent analysis and visualization were performed in R using a variety of tools and packages. Correlation plots were generated using base R and the corrplot package v0.92 (WEI AND SIMKO 2021). Principal components were analyzed and visualized with PCAtools v2.2.0 (BLIGHE AND LUN 2022). Heatmaps, including clustering analysis performed by the hclust method (from base R {stats} package), were generated by pheatmap v1.0.12 (KOLDE 2019).

Gene Ontology Analysis was carried out using the GOrilla tool for genes that were significantly (*P*-value <0.05) up-regulated or down-regulated by more than two-fold in *Pyk* mutants (EDEN *et al*. 2009). Tissue enrichment analysis was conducted using default settings for the “Preferred tissue (modEncode RNA_seq)” tool in PANGEA version 1.1 beta (HU *et al*. 2023).

## Data Availability

All strains and reagents are available upon request. All targeted metabolomics data described herein are included in Tables S2 and S3. Processed RNA-seq data is presented in Table S4 and available in NCBI Gene Expression Omnibus (GEO).

## RESULTS

### The *Drosophila* gene *Pyk* encodes the Pyk enzyme orthologous to human PKs

The *Drosophila* genome harbors six genes predicted to encode pyruvate kinases (GARAPATI *et al*. 2019; GRAMATES *et al*. 2022b). To better understand the relationships between the *Drosophila* Pyk paralogs and Pyk orthologs in other organisms, we compared their pairwise protein sequence identities and generated a phylogenetic gene tree to infer their evolutionary history (Figure 1, Table S1). Species examined included human and mouse, *Aedes aegypti, Caenorhabditis elegans*, *Saccharomyces cerevisiae*, and *Arabidopsis thaliana,* with *Escherichia coli* as the outgroup for tree rooting (Table S1). The pairwise sequence comparisons indicate that *Drosophila* Pyk shares strongest identity with that of the *Aedes aegypti* protein PK (XP_001649983.1), followed by those of its mammalian orthologs, the worm paralogs, and fly paralog CG7069 (Figure 1A).

**Figure 1.**
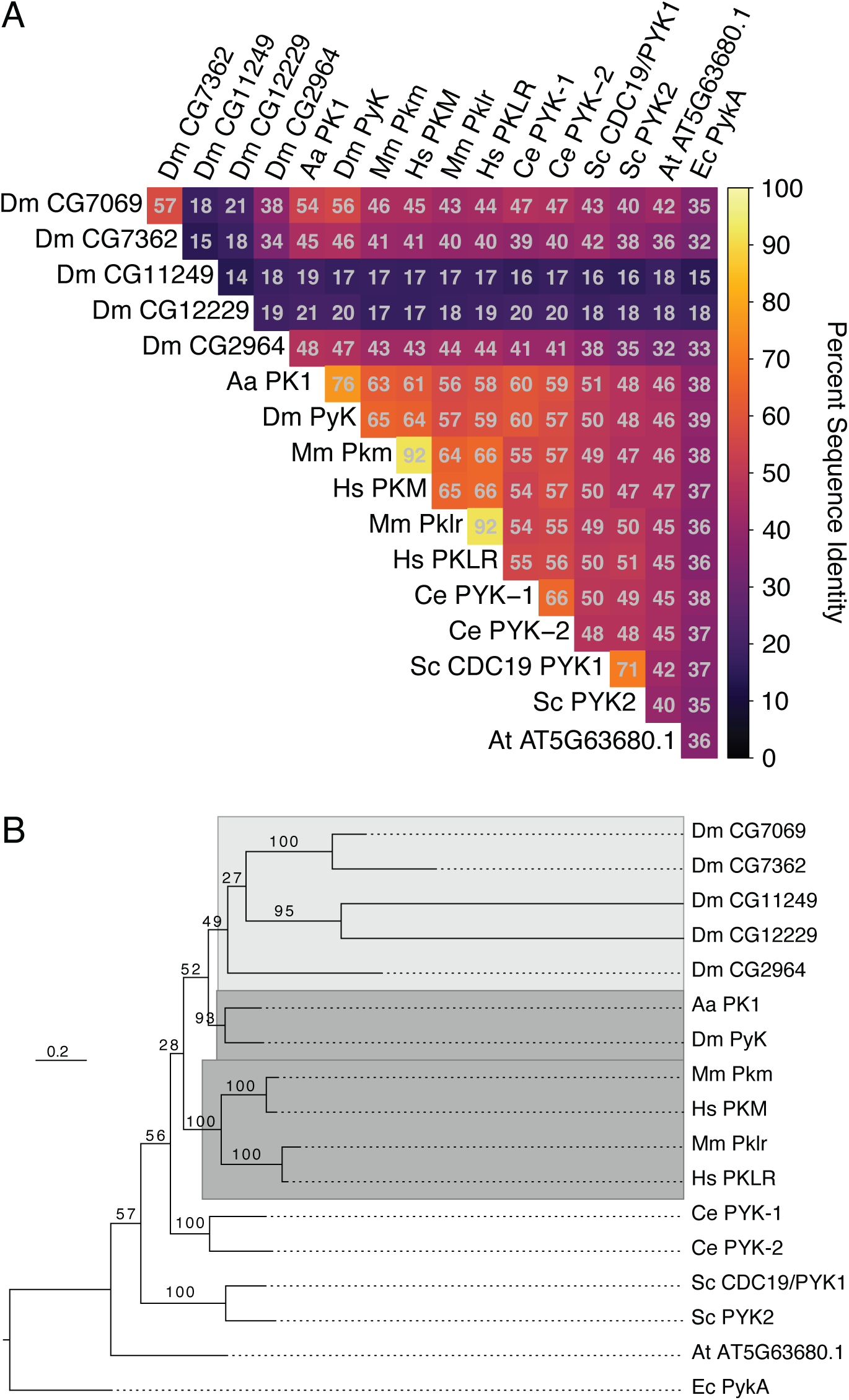
Sequence identity and phylogeny of Pyk homologs. (A) Heatmap of pairwise sequence identities extracted from the ensemble MSA. (B) Phylogenetic maximum likelihood gene tree. Ultrafast bootstrap support is shown on internal branches. The dark grey highlighted clades include the canonical *Drosophila melanogaster*, *Aedes aegypti*, and human/mouse orthologs, while the light grey region encompasses the divergent fly paralogs. The tree is rooted to *E*. c*oli* homolog PykA as the outgroup. Species abbreviations: Aa, *A*. *aegypti; Dm*, *D. melanogaster*; Hs, *H. sapiens*; Mm, *M. musculus*; Ce, *C. elegans*; Sc, *S. cerevisiae*; At, *A. thaliana*; Ec, *E. coli*. See Supplemental Table 1 for a list of isoforms used in this analysis.

The other putative paralogs show identity to a lesser extent, with two of the *Drosophila* proteins (CG11249 and CG12229) displaying less than 20% identity with all other analyzed Pyk orthologs (Figure 1A). The sequence similarities, allowing for conservative residue substitutions, show a similar pattern (Figure S2). These results indicate that, at the sequence level, *Drosophila* Pyk is as or more similar to the yeast orthologs as it is to 4 of the 5 other putative fly paralogs.

Sequence distances may not correspond to actual evolutionary or functional relationships, so we next sought to reconstruct the Pyk gene tree using a maximum likelihood model of protein evolution. This phylogeny indicates that the *Drosophila* Pyk is sister to its *Aedes aegypti* ortholog (Figure 1B), which composes a larger clade with the more distantly related *Drosophila* paralogs. This insect Pyk/PK clade is in turn sister to that of the mammalian orthologs, which share a common ancestor. We note strong ultrafast bootstrap support for the insect and mammalian ortholog subclades, the yeast and worm paralogs, and two pairs of the fly paralogs, with weak support for the fly paralog internal branches and the relationships between species (Figure 1B).

*Aedes* PK1 was previously demonstrated to exhibit unique allosteric regulation when compared with mammalian enzymes (PETCHAMPAI *et al*. 2019). Given the high sequence similarity and close evolutionary relationship between these two Dipteran proteins, our findings suggest that *Drosophila* Pyk functions in a similar manner to that of mosquito PK. Overall, these results support the hypothesis that the fly gene *Pyk* encodes the canonical glycolytic enzyme, which is consistent with previous studies demonstrating that *Pyk* is the most widely expressed member of the *Drosophila melanogaster* Pyk gene class (RUST AND COLLIER 1985; GRAVELEY *et al*. 2011).

### *Pyk* mutants die during the second larval instar

To determine how loss of Pyk activity affects *Drosophila* development, we used CRISPR/Cas9 to generate two small deletions in the *Pyk* locus, *Pyk^23^* and *Pyk^31^*, both of which induce frameshifts in the first coding exon and are predicted to be null alleles (Figure 2A). When these two *Pyk* mutant alleles were placed in trans, the resulting *Pyk^23/31^* mutants survived embryogenesis but died during the mid-L2 stage (Figure 2B). Moreover, when compared with *Pyk^23/+^* heterozygous controls, *Pyk* mutants displayed a significant decrease in both body mass and triglyceride stores (Figure 2C and 3A, B), thus demonstrating that loss of Pyk activity significantly disrupts larval development. We would note, however, that although *Pyk* mutants accumulate far less triglycerides (TAG) than controls, the density of lipid droplets within the fat body as determined by Nile Red staining (Figure 3A,B) remains similar to controls. Overall, these findings reveal that *Pyk* mutant larvae exhibit many of the same phenotypes observed in *dERR* and *Pfk* mutants, which also exhibit severe disruptions of glycolytic metabolism (RUST AND COLLIER 1985; TENNESSEN *et al*. 2011).

**Figure 2.**
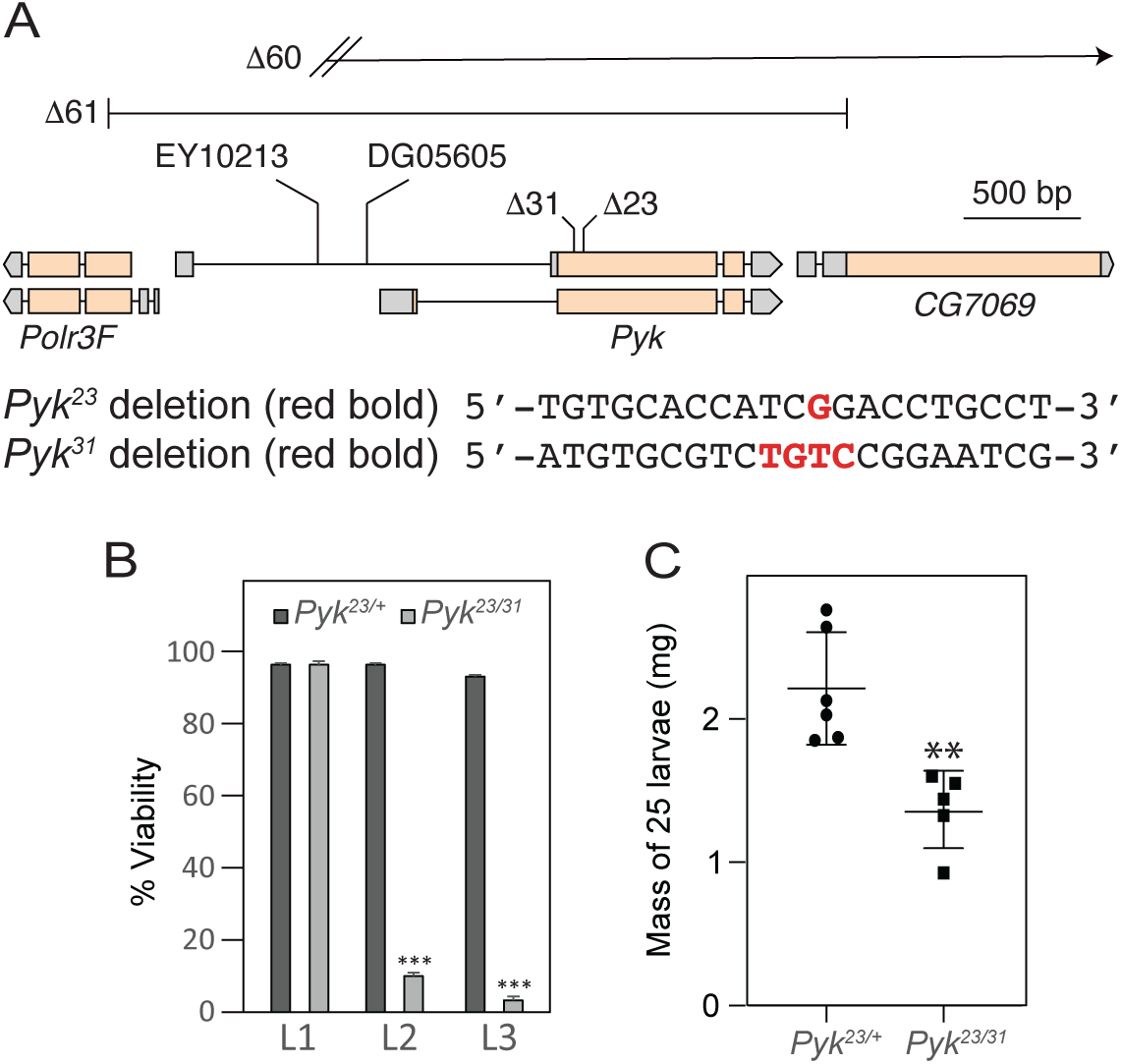
Deletions in the *Pyk* coding region induce larval lethality and growth defects. (A) An illustration of the *Drosophila Pyk* locus and surrounding genes, as well as the *Pyk* mutant alleles. The *Pyk^23^* and *Pyk^31^* alleles were generated using a CRISPR/Cas9-based strategy and induce small deletions that result in frameshifts. The *Pyk^60^* and *Pyk^61^* alleles represent deletions that remove the entire PYK coding region as well as portions of neighboring genes. See Figure S1 for analysis of *Pol3f* and *CG 7069* gene expression in trans-heterozygous *Pyk* mutant backgrounds. (B) A histogram illustrating the effects of *Pyk* mutations on larval viability. Note that nearly all *Pyk^23/31^* trans-heterozygous mutant animals die during the second larval instar – phenocopying *dERR* mutants. *P <* 0.001 (C) *Pyk^23/31^* mutants are significantly smaller than heterozygous controls. n=6 samples containing 25 mid-L2 larvae. \*\**P <* 0.01. \*\*\**P <* 0.001. *P*-values calculated using a Mann Whitney test.

**Figure 3.**
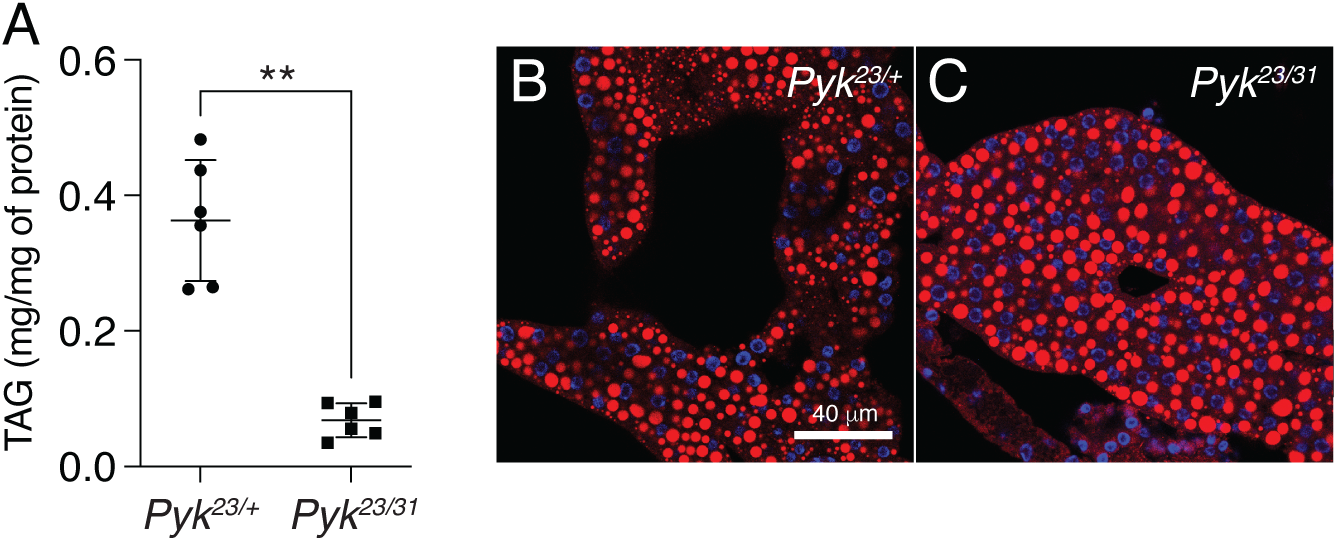
*Pyk* mutants exhibit decreased TAG levels. TAG levels were assayed during L2 development in *Pyk* mutants and heterozygous controls. (A) *Pyk^23/31^* mutants exhibit significantly decreased triglycerides (TAG) as compared with *Pyk^21/+^* controls. Data are normalized to soluble protein. n=6 biological replicates containing 25 mid-L2 larvae per sample. \*\**P <* 0.01. *P*-values calculated using a Mann Whitney test. (B,C) Nile Red was used to stain lipid droplets in L2 fat bodies of both (B) *Pyk^21/+^* controls and (C) *Pyk^23/31^* mutants. Scale bar in (B) applies to (C).

### Metabolomic analysis of *Pyk* mutants

To better understand how loss of *Pyk* alters larval metabolism, we used a targeted metabolomics approach to compare *Pyk^23/31^* mutants with *Pyk^23/+^* heterozygous controls during the L2 larval stage (Table S2). Partial Least Squares Discriminant Analysis (PLS-DA) of the resulting data revealed that control and mutant samples clustered in distinct groups (Figure S3). Subsequent analysis of the data revealed increased levels of the metabolites immediately upstream of Pyk, 2/3-phosphoglycerate and phosphoenolpyruvate, and a decrease in lactate, and 2-hydroxyglutarate, and most amino acids that can be used as anaplerotic fuel in the citric acid cycle (Figure 4A-B, Table S2). Overall, these changes are indicative of a block in glycolysis at the reaction catalyzed by Pyk.

**Figure 4.**
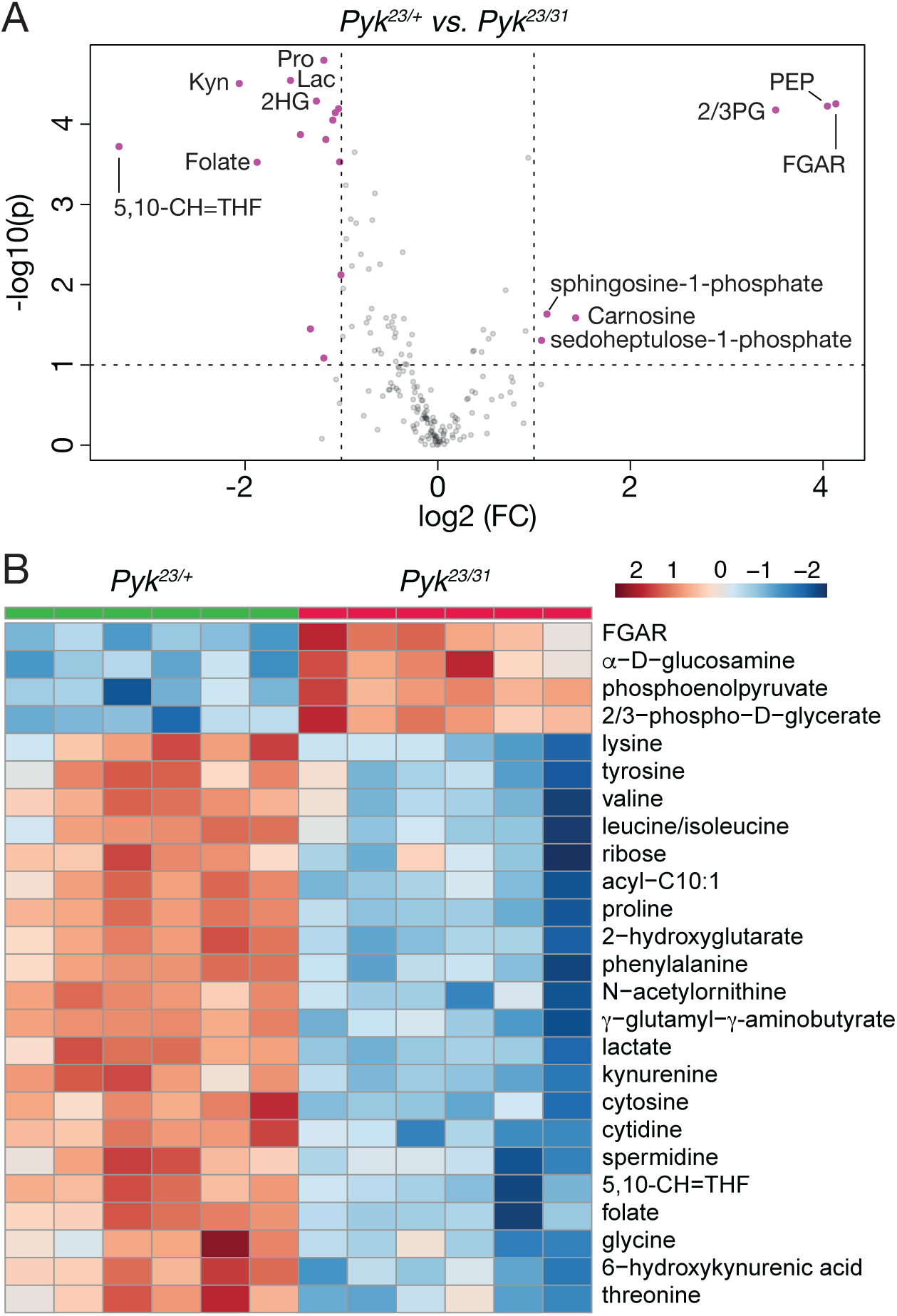
Targeted metabolomic analysis of *Pyk* mutant larvae. *Pyk^23/31^* mutant larvae and *Pyk^21/+^* controls were analyzed using a targeted GC-MS approach. (A) Differences in metabolite abundance between control and mutant samples are represented as a volcano plot. Dashed vertical line represents a log2 fold change (FC) of > 1.5. Dashed horizontal line represents p < 0.01. (B) A heat map illustrating the top 25 most significantly altered compounds in *Pyk^23/31^* mutants as compared with *Pyk^21/+^* controls. Both figures were generated using Metaboanalyst 5.0, as described in the methods. Abbreviated compounds are 2/3-phosphoglycerate (2/3PG), phosphoenolpyruvate (PEP), 2-hydroxyglutarate (2HG), 3-hydroxybutyrate (2HB), 5,10-Methenyltetrahydrofolate (5,10-CH=THF), alanine (Ala), Asparagine (Asn), Glutamic acid (Glu), Glutamine (Gln), 5’−Phosphoribosyl−N-formylglycinamide (FGAR), proline (Pro), trehalose (Tre).

While the metabolite changes observed in our initial metabolomics dataset were largely expected, we were surprised to find that steady state pyruvate levels were unchanged in *Pyk* mutants (Table S2). To verify that pyruvate levels remained constant even in the absence of Pyk, we analyzed a second series of *Pyk* mutants using a different methodology in an independent metabolomics core facility. In this case, a trans-heterozygous combination of two *Pyk* deletion alleles, *Pyk^60/61^*, were compared with a genetically-matched control strain that was homozygous for a precise excision allele (Table S3; see Methods for a description of the alleles used). Although these deletions also removed portions of *CG7069* (Figure 2A), which encodes another member of the Pyk family, this gene is not expressed during most of larval development, and thus our analysis likely reflects phenotypes caused by the loss of the *Pyk* locus. Similar to the metabolomic profile of *Pyk^23/31^* mutants, *Pyk^60/61^* mutant L2 larvae exhibited increased levels of glycolytic metabolites immediately upstream of pyruvate kinase activity (2-phosphoglycerate, 3-phosphoglycerate and phosphoenolpyruvate), and decreased levels of the downstream metabolites lactate, 2HG, and several amino acids (Figure 5A,B; Table S3). Moreover, these *Pyk* mutant larval samples also displayed normal pyruvate levels (Figure 5A), suggesting that steady state levels of larval pyruvate are maintained by compensatory changes in other metabolic pathways.

**Figure 5.**
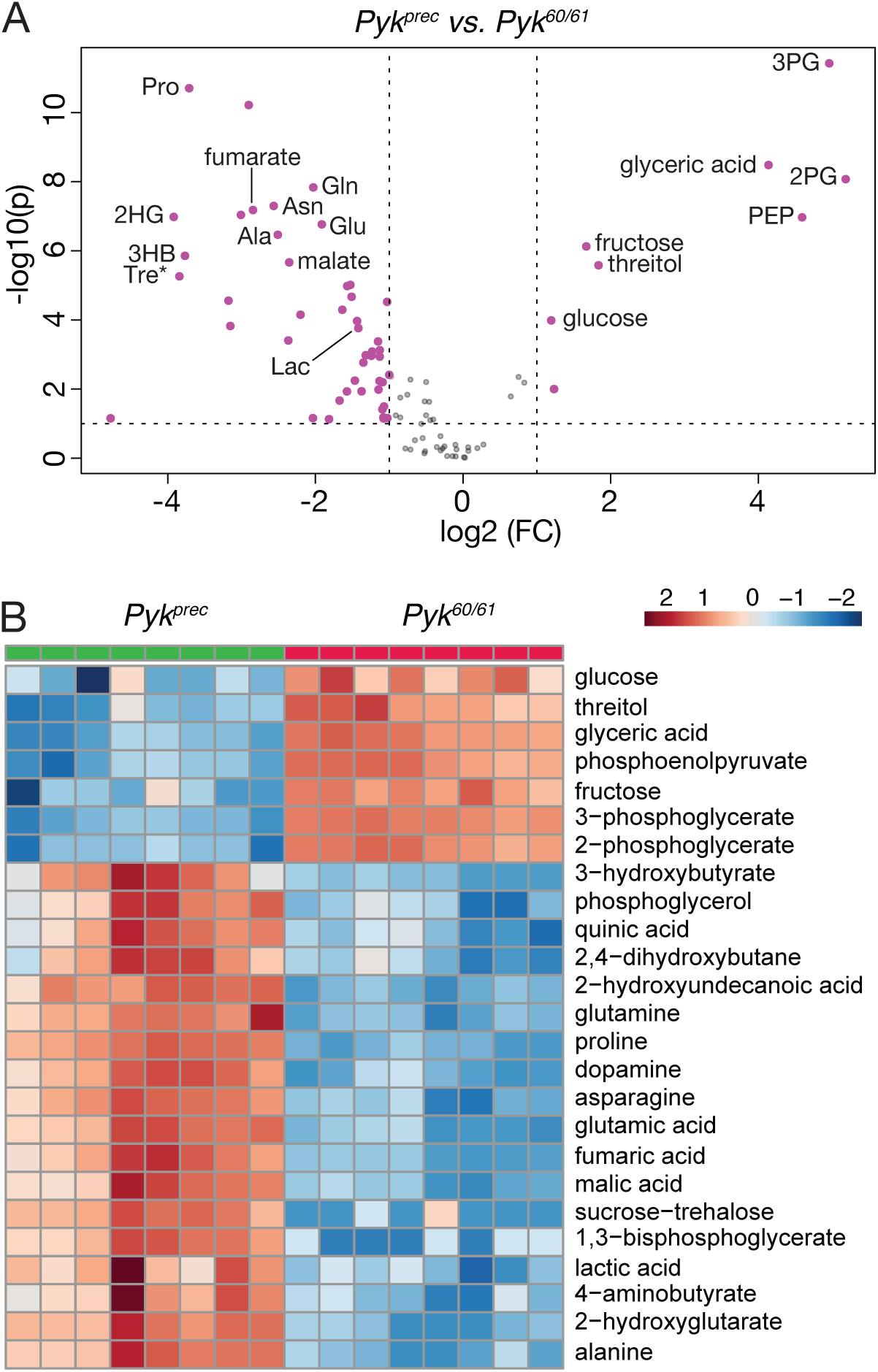
Targeted metabolomic analysis of *Pyk* mutant larvae. *Pyk^60/61^* mutant larvae and precise excision controls (*Pyk^Prec^*) were analyzed using a targeted GC-MS approach. (A) Differences in metabolite abundance between control and mutant samples are represented as a volcano plot. Dashed vertical line represents a fold change (FC) of > 1.5. Dashed horizontal line represents *P* < 0.01. (B) A heat map illustrating the top 25 most significantly altered compounds in *Pyk^60/61^* mutants as compared with *Pyk^Prec^* controls. Both figures were generated using Metaboanalyst 5.0, as described in the methods. Abbreviated compounds are 3-phosphoglycerate (3PG), 2-phosphoglycerate (2PG), phosphoenolpyruvate (PEP), 2-hydroxyglutarate (2HG), 3-hydroxybutyrate (2HB), alanine (Ala), Asparagine (Asn), Glutamic acid (Glu), Glutamine (Gln), 5’−Phosphoribosyl−N-formylglycinamide (FGAR), proline (Pro), trehalose (Tre).

As an extension of our metabolomic studies, we compared our datasets to identify those metabolites that were similarly altered in both *Pyk* mutant backgrounds. While we would note that this comparative analysis is somewhat limited in that we used two different targeted metabolomics protocols that were optimized to detect different subsets of metabolites, molecules that appear in both analyses highlight metabolic pathways that are significantly altered by loss of Pyk activity in different genetic backgrounds. Despite the many metabolic changes observed in each individual dataset, only a handful of metabolites were common among the two experiments: 2/3-phosphoglycerate, phosphoenolpyruvate, lactate, 2-hydroxyglutarate, and proline (Figure 6A-C). While the changes in 2/3-phosphoglycerate, phosphoenolpyruvate, and lactate were to be expected, the decrease in 2HG and proline present an interesting model for how metabolic flux could be rewired in the absence of Pyk activity. The decrease in 2HG likely represents L-2HG, which is abundant in larvae (LI *et al*. 2017; MAHMOUDZADEH *et al*. 2020). Since L-2HG is synthesized from α-ketoglutarate by LDH and is sensitive to changes in lactate production (LI *et al*. 2017; LI *et al*. 2018), we hypothesize that the decreased abundance of 2HG represents a smaller lactate pool combined with a potential decrease in TCA cycle flux. Meanwhile, the decreased proline levels present in *Pyk* mutants are quite informative because this amino acid is commonly used by insects as an anaplerotic fuel for the TCA cycle (BURSELL 1981). Based on this observation, future studies should determine if *Pyk* mutants are using proline to sustain both pyruvate levels and early larval growth.

**Figure 6.**
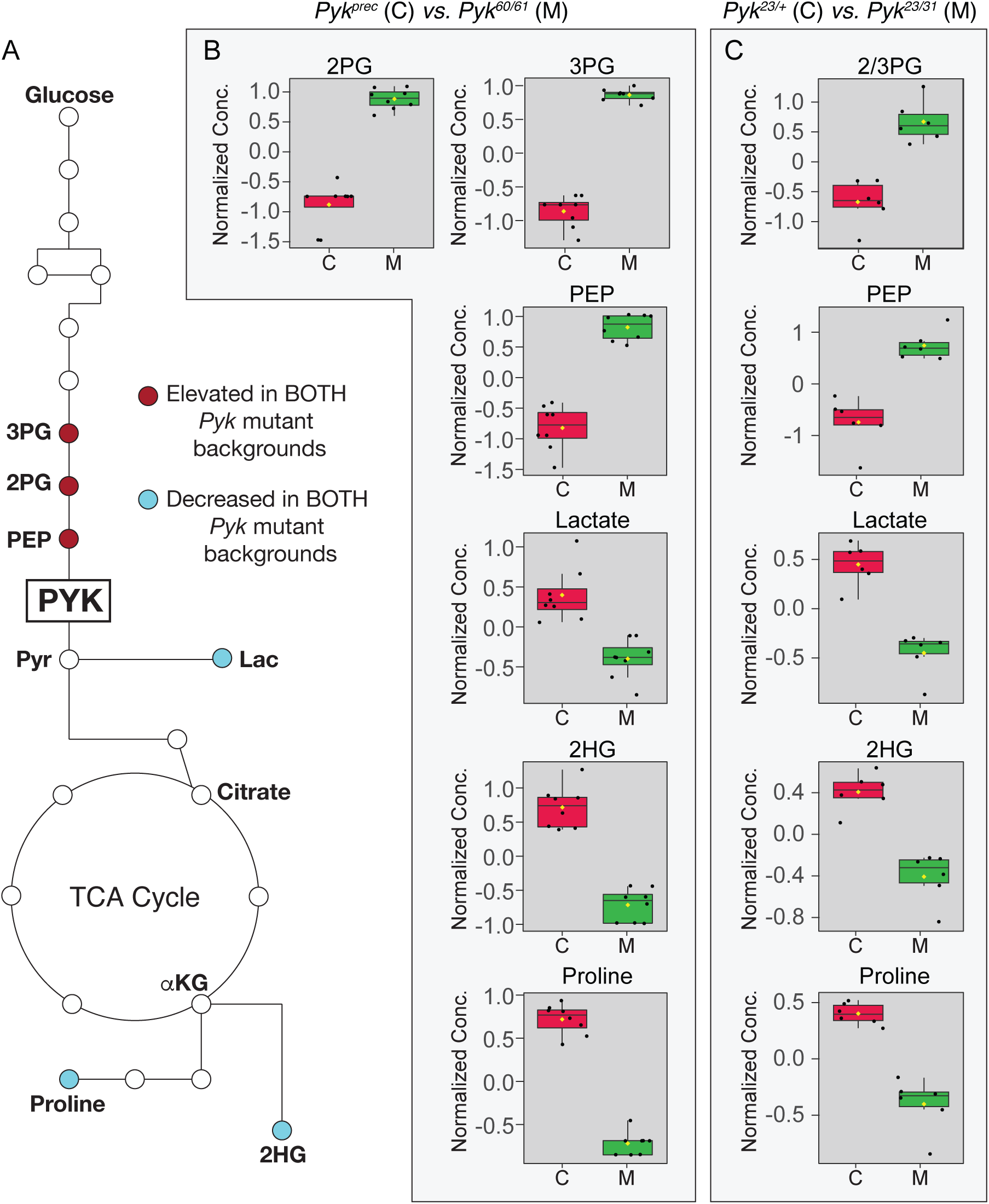
A limited set of metabolites are significantly altered in both *Pyk* mutant backgrounds. (A) An illustration of glycolysis and the TCA cycle with metabolites that are significantly altered in both *Pyk* mutant backgrounds highlighted in dark red (up-regulated) and light blue (down-regulated). (B,C) Boxplots illustrating the change in abundance of metabolites that are significantly altered in both (B) *Pyk^60/61^* and (C) *Pyk^23/31^* mutant backgrounds. All boxplots were generated using Metaboanalyst 5.0, as described in the methods. Black dots represent individual samples, the horizontal bar in the middle represents the median and the yellow diamond represents the mean concentration. For all boxplots, the metabolite fold-change was >2 fold and *P <* 0.01. Abbreviated compounds are 3-phosphoglycerate (3PG), 2-phosphoglycerate (2PG), phosphoenolpyruvate (PEP), 2-hydroxyglutarate (2HG).

### Gene Expression Analysis of *Pyk* mutants

Our metabolomic studies suggest that loss of Pyk activity induces compensatory changes in other metabolic pathways that allow the larvae to survive early development. To better understand if this metabolic rewiring is recapitulated at the level of gene expression, we used RNA-seq to compare the expression profiles of *Pyk^23/31^*mutant larvae with *Pyk^23/+^* heterozygous controls (Table S4). This analysis revealed significant differences in the gene expression between mutant and control samples (Figure S5A,B), with 755 genes up-regulated and 214 genes downregulated more than two-fold in the *Pyk* mutant dataset (adjusted *P* value <= 0.05; Table S4, Figure 7). Notably, the expression of the *Pyk* gene itself was down-regulated 8.3-fold when compared with the control strain, thus demonstrating the severity of these deletion alleles (Table S4, Figure 7). The data also revealed a significant upregulation of the gene *Thor* (FBgn0261560), the fly homolog of 4E-BP (BERNAL AND KIMBRELL 2000), which functions as a metabolic brake that dampens mRNA translation (HAY AND SONENBERG 2004; TELEMAN *et al*. 2005). The significant upregulation of *Thor* transcript levels hints at one possible mechanism by which *Pyk* mutants both rewire larval metabolism and slow growth in response to a major metabolic insult.

**Figure 7.**
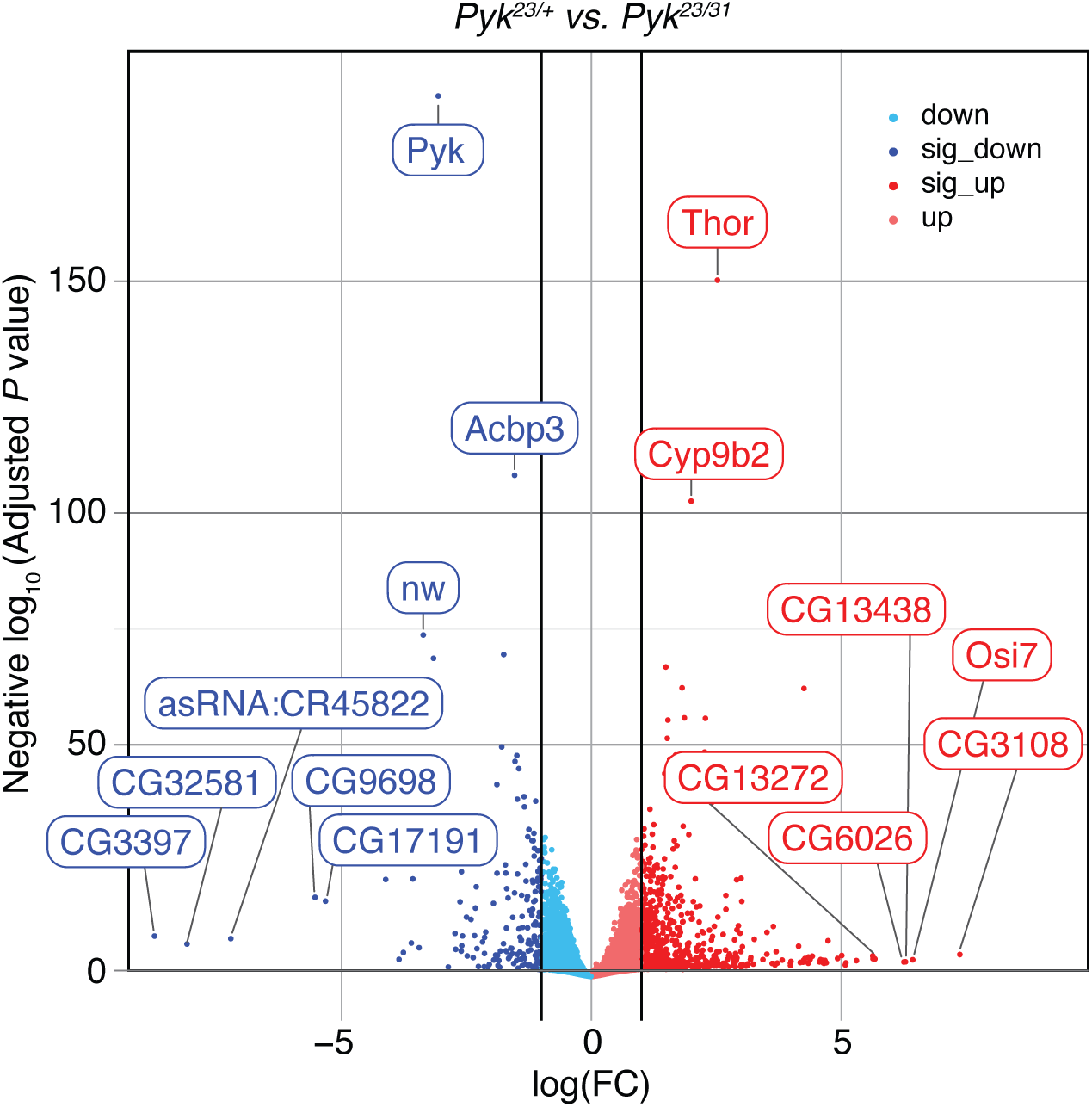
RNA-seq analysis of *Pyk^23/31^* mutants as compared with *Pyk^23/+^* controls. A volcano plot illustrating gene expression changes in *Pyk^23/31^* mutants as compared with *Pyk^23/+^* controls. Black vertical lines indicate absolute log fold change (LFC) ≥ 1. Black horizontal line represents adjusted *P* value ≤ 0.05. The top 15 most significantly changed genes are labeled, with all significant genes (abs(LFC) ≥ 1 and adj *p* ≤ 0.05) shaded darker.

To further analyze the *Pyk* mutant RNA-seq data, we used Gene Ontology (GO) to better define the biological processes most affected by loss of Pyk activity (Figure 8A,B and Table S5). Our analysis revealed that *Pyk* mutant samples exhibited a significant enrichment in GO categories associated with the digestion and metabolism of chitin and glucosamines, lipids, and proteins (Figure 8A). Notably, analysis of the RNA-seq data set using the “Preferred tissue” tool in PANGEA indicated that the digestive system displays the largest number of significantly up-regulated genes in *Pyk* mutants (Table S6) (HU *et al*. 2023), with many of the those genes being involved in lipid and protein metabolism (Figure 8C). For example, lipid metabolism was the second-most enriched GO category, and; most of the significantly up-regulated genes that fit within this GO category encode intestinal lipases. Similarly, *Pyk* mutants exhibit significant increases in genes that encode proteases (Figure 8A, C), many of which represent digestive proteases predicted to be expressed within the intestine. Overall, our results suggest that the metabolism of *Pyk* mutants attempts to compensate for its loss by increasing expression of genes involved in the breakdown of macromolecules within the digestive tract, perhaps as a means of increasing the uptake of dietary nutrients.

**Figure 8.**
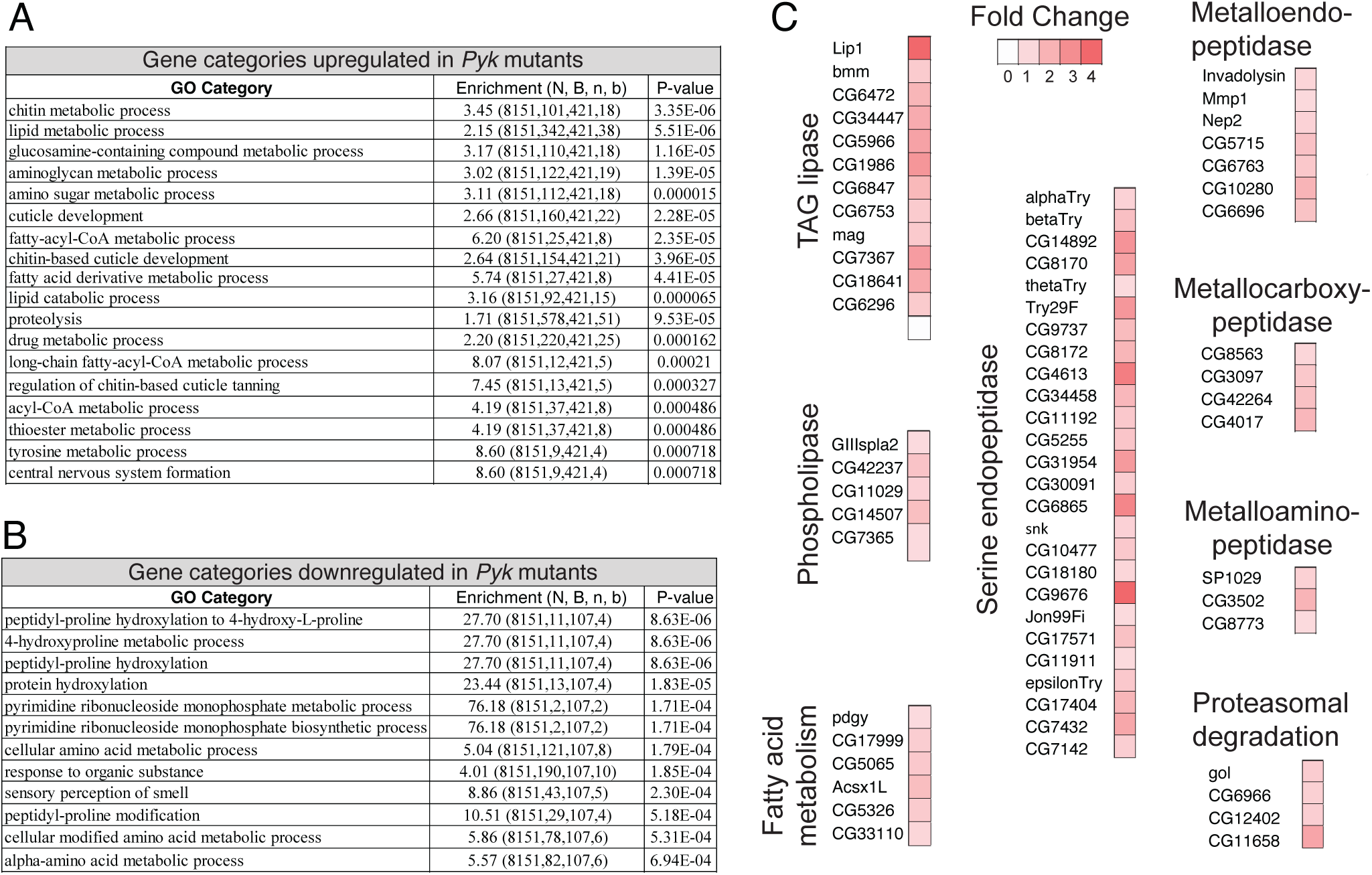
Gene Ontology (GO) enrichment analysis of genes that are significantly misregulated in *Pyk^23/31^* mutants. GOrilla was used to determine GO enrichment among those genes that were either (A) significantly up-regulated or (B) significantly down-regulated in *Pyk^23/31^* mutants when compared to the *Pyk^23/+^* control. (C) Significantly up-regulated genes in the GO categories “lipid metabolic process” and “proteolysis” are grouped by enzymatic function, with the increased fold-change in gene expression represented by a color gradient.

## DISCUSSION

Here we demonstrate that *Pyk* mutants exhibit a larval lethal phenotype with defects in larval growth and a severe block in glycolysis. While the phenotypic consequences of mutating *Pyk* are largely expected, our study contains several important findings for both the *Drosophila* and biomedical communities. First, we demonstrate that while the *Drosophila* genome encodes several pyruvate kinase orthologs, the gene *Pyk* encodes the canonical enzyme most analogous to Pyk in other organisms. In fact, *Drosophila* Pyk is more similar to the two yeast orthologs than to the other five pyruvate kinase orthologs present in the fly. Future studies should more closely examine these paralogous enzymes and determine why *Drosophila melanogaster* evolution produced such a specialized set of pyruvate kinases.

A second key finding from our sequence analysis is that *Drosophila* Pyk exhibits nearly equivalent sequence similarity to both the mammalian PKLR and PKM orthologs. This observation is important because PLKR and PKM are connected with distinct disease phenotypes. Thus, future studies in the fly should be cautious when making comparisons with disease states that are unique to either of the human enzymes. Instead, the *Pyk* mutants should be used as a tool to understand how cells and tissues respond to a general deficiency in Pyk activity, similar to previous studies that manipulated *Drosophila* Pyk activity in imaginal disc development and neurons/glia (TIXIER *et al*. 2013; VOLKENHOFF *et al*. 2015; WU *et al*. 2018; SPANNL *et al*. 2020). One exciting possibility would be to use our newly described mutants as a background for expressing disease causing variants of the human enzymes. After all, pyruvate kinase deficiency is the most common enzymatic disorder detected in newborns, and moreover, a wide variety of disease-causing PKLR variants have been identified (DOLAN *et al*. 2002; GRACE *et al*. 2015; FATTIZZO *et al*. 2022; LUKE *et al*. 2023). Expression of the wild-type and mutated enzyme variants in the fly would allow for a rapid genetic analysis of how PKLR mutations alter cellular metabolism while also providing an *in vivo* platform to test potential therapeutics.

Our studies of the *Pyk* mutant also reveal how a block in glycolysis alters systemic metabolism in unexpected ways. Previous studies in the fly have examined how disruption of glycolytic metabolism affects both the metabolome and transcriptome. For example, the *Drosophila* Estrogen-Related Receptor (dERR) is a central regulator of glycolytic metabolism throughout the fly lifecycle (TENNESSEN *et al*. 2011; BEEBE *et al*. 2020). Our studies reveal that the developmental defects exhibited by *Pyk* mutants largely phenocopy those observed in *dERR* mutants although the *Pyk* and *dERR* mutant metabolomic and transcriptomic profiles differ significantly. Unlike *dERR* mutants, which display a buildup of sugars due to the down-regulated expression of nearly every glycolytic enzyme (TENNESSEN *et al*. 2011; BEEBE *et al*. 2020), *Pyk* mutants instead accumulate three glycolytic metabolites immediately upstream of the Pyk-catalyzed reaction. Thus, our studies of *Pyk* highlight how the *dERR* mutant metabolic profile represents a broad disruption of carbohydrate metabolism when compared with the loss of a single *dERR* target gene.

While most aspects of the *Pyk* mutant metabolomics profile were expected based on the known role of Pyk, several interesting observations emerged. First, steady state levels of pyruvate were unchanged in *Pyk* mutants as compared with controls. Since the other metabolites in glycolysis are changed in a manner that is indicative of a loss of Pyk activity, and both lactate and TCA cycle intermediates exhibit a severe decrease, flux into and out of the larval pyruvate pool must be severely compromised. Future studies should explore this possibility using stable-isotope tracers and further examine how larval metabolism maintains steady state pyruvate levels even under severe metabolic stress.

Our analysis also raises questions about how *Pyk* mutants, as well as *dERR* mutants, survive until the mid-L2 stage. Glycolysis is severely disrupted in both mutant backgrounds, yet these animals continue to develop for almost three days when raised under standard culture conditions. While this impressive resilience might be partially explained by maternal loading of Pyk and other enzymes, both metabolomics and transcriptomics analysis suggest that *Pyk* mutants switch fuel sources in an attempt to maintain developmental progress. Notably, *Pyk* mutants exhibit a significant decrease in the fermentation products lactate and 2HG, as well as a decrease in the amino acid proline. All three changes signal a major reorganization of larval metabolism, as lactate and L-2HG are present at high levels within normal larvae and represent pools of carbon that are not immediately used for energy production and growth (LI *et al*. 2017). Meanwhile, proline is commonly used for energy production in insects (TEULIER *et al*. 2016). Thus the depletion of lactate, 2HG, and proline steady state pools suggests that *Pyk* mutant metabolism has become reliant on alternative energy sources, such as proline, and lacks adequate resources to ferment pyruvate and α-ketoglutarate into lactate and L-2HG, respectively. This hypothesis is also supported by RNA-seq analysis, which uncovered a significant up-regulation of enzymes involved in the digestion and metabolism of complex lipids and proteins. Overall, further studies of *Pyk* mutants are warranted to understand how these metabolic adaptations are induced and regulated.

Our results also revealed that *Pyk* mutants exhibit more severe developmental phenotypes when compared with *Mpc1* mutants, which are unable to properly transport pyruvate into the mitochondria (BRICKER *et al*. 2012). Considering that Pyk and Mpc1 both directly affect pyruvate metabolism, one might predict that mutations in these genes would result in similar phenotypes. Instead, *Mpc1* mutants are viable when raised on standard *Drosophila* media and only exhibit a lethal phenotype when placed on a sugar-only diet (Bricker et al., 2012). Moreover, the metabolomics profile of *Mpc1* and *Pyk* mutants display significant differences. Notably, serine levels are significantly increased in *Mpc1* mutants – an observation that would be consistent with an uncoupling of glycolysis and the TCA cycle causing a rerouting of glycolytic intermediates into the serine/glycine biosynthetic pathway (Bricker et al.,2012). In contrast, *Pyk* mutants exhibit no increase in serine production despite harboring very high levels of 3-phosphoglycerate, which serves as the precursor for serine biosynthesis (YE *et al*. 2012). This difference is surprising because studies of PKM2 in cancer cells suggests that decreased Pyk activity induces shunting of glycolytic intermediates into serine production (YE *et al*. 2012).

One explanation for the dramatic phenotypic differences between *Pyk* and *Mpc1* mutants stems from differences in lactate metabolism. Unlike *Pyk* mutants, *Mpc1* mutants accumulate excess lactate. Thus, *Mpc1* mutants can generate two more ATPs from each glucose molecule when compared with *Pyk* mutants, while also regenerating ATP independent of mitochondrial activity. Future studies should test this model by examining if *Mpc1;Ldh* double mutant exhibits metabolic and developmental phenotypes that more closely mimic those observed in *Pyk* mutants.

In conclusion, we have here described a metabolomics profile of *Drosophila* larvae lacking Pyk activity. While the resulting changes in central carbon metabolism are largely expected, such an analysis of fly metabolism is essential as *Drosophila* geneticists increasingly study the role of metabolism across a wide range of contexts. Moreover, *Drosophila* metabolism is highly adaptable and compensates for metabolic insult – as seen here by depletion of the anaplerotic amino acids proline, glutamate, and glutamine. A careful exploration of these metabolic networks will be essential for studying metabolism in the context of *Drosophila* growth, development, and models of human disease.

## ACKNOWLEDGEMENTS

Thanks to the Bloomington *Drosophila* Stock Center (NIH P40OD018537) for providing fly stocks, the *Drosophila* Genomics Resource Center (NIH 2P40OD010949) for genomic reagents, and Flybase (NIH 5U41HG000739). Metabolomics analysis performed at the Metabolomics Core Facility at the University of Utah is supported by spectrometry equipment obtained through NCRR Shared Instrumentation Grants 1S10OD016232-01, 1S10OD018210-01A1 and 1S10OD021505-01. Sequencing was performed at Indiana University’s Center for Genomics and Bioinformatics. All computation was performed on Indiana University’s Carbonate and Research Desktop HPC clusters. We thank members of the Matt Hahn lab at IU for helpful guidance on the phylogeny. J.M.T. is supported by the National Institute of General Medical Sciences of the National Institutes of Health under a R35 Maximizing Investigators’ Research Award (MIRA; 1R35GM119557).

## SUPPLEMENTAL FIGURE LEGENDS

Figure S1. ***Pyk* mRNA transcript levels are significantly reduced in *Pyk* mutant larvae.** Total RNA from stage *w^1118^; Pyk^prec^* control larvae and *w^1118^; Pyk^60/61^* mutant larvae were analyzed by northern blot hybridization to detect transcripts encoding Pyk, CG18596, and Polr3F. Hybridization to detect *rp49* mRNA is included as a loading control.

Figure S2. **A comparison of sequence similarities between Pyk homologs.** A heatmap of pairwise sequence identities extracted from the ensemble MSA. Species abbreviations: Aa, *A. aegypti; Dm*, *D. melanogaster*; Hs, *H. sapiens*; Mm, *M. musculus*; Ce, *C. elegans*; Sc, *S. cerevisiae*; At, *A. thaliana*; Ec, *E. coli.* See Supplemental Table 1 for a list of isoforms used in this analysis.

Figure S3. A comparison of the metabolomic data from *Pyk^23/31^* mutant and *Pyk^21/+^*control samples using principal component (PC) analysis. Targeted metabolomics data from Table S1 was analyzed using principal component analysis. Analysis was conducted using Metaboanalyst 5.0.

Figure S4. A comparison of the metabolomic data from *Pyk^60/61^* mutant and *Pyk^prec^*control samples using principal component (PC) analysis. Targeted metabolomics data from Table S2 was analyzed using principal component analysis. Analysis was conducted using Metaboanalyst 5.0.

Figure S5. **RNA-seq Principal Component Analysis and Correlations of Experimental Replicates.** (A) PCA biplot of principal component 1 (PC1) versus PC2 for the top 5,000 genes by variance, with replicates labeled. (B) Spearman correlation plot of the top 5,000 genes by variance after filtering out genes with either low/no expression.

## SUPPLEMENTAL TABLE LEGENDS

**Table S1.** Protein isoform information for the Pyk orthologs used in Figure 1 and Figure S2.

**Table S2.** Metabolomic analysis of *Pyk^23/31^* mutants as compared with *Pyk^23/+^* controls.

**Table S3.** Metabolomic analysis of imprecise excision *Pyk* mutants as compared with precise excision controls.

**Table S4.** RNA-seq results comparing gene expression between Pyk23/31 mutants and *Pyk*^23/+^.

**Table S5.** GO analysis of significantly mis-regulated genes from the RNAseq data represented in Table S4 using GOrilla.

**Table S6.** Analysis of RNA-seq data from Table S4 using the Preferred tissue (modEncode RNA_seq) tool in PANGEA.

