## Supplementary material for "Metabolomic analysis of *Drosophila melanogaster* larvae lacking Pyruvate kinase": Figure S1

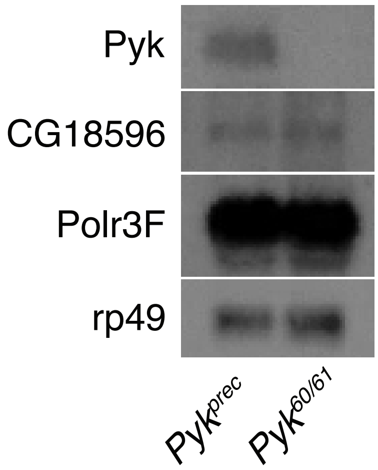


**Figure S1. *Pyk* mRNA transcript levels are significantly reduced in *Pyk* mutant larvae.** Total RNA from stage $w^{1118}$; $Pyk^{prec}$ control larvae and $w^{1118}$; $Pyk^{60/61}$ mutant larvae were analyzed by northern blot hybridization to detect transcripts encoding Pyk, CG18596, and Polr3F. Hybridization to detect *rp49* mRNA is included as a loading control.
