## Supplementary material for "Metabolomic analysis of *Drosophila melanogaster* larvae lacking Pyruvate kinase": Figure S2

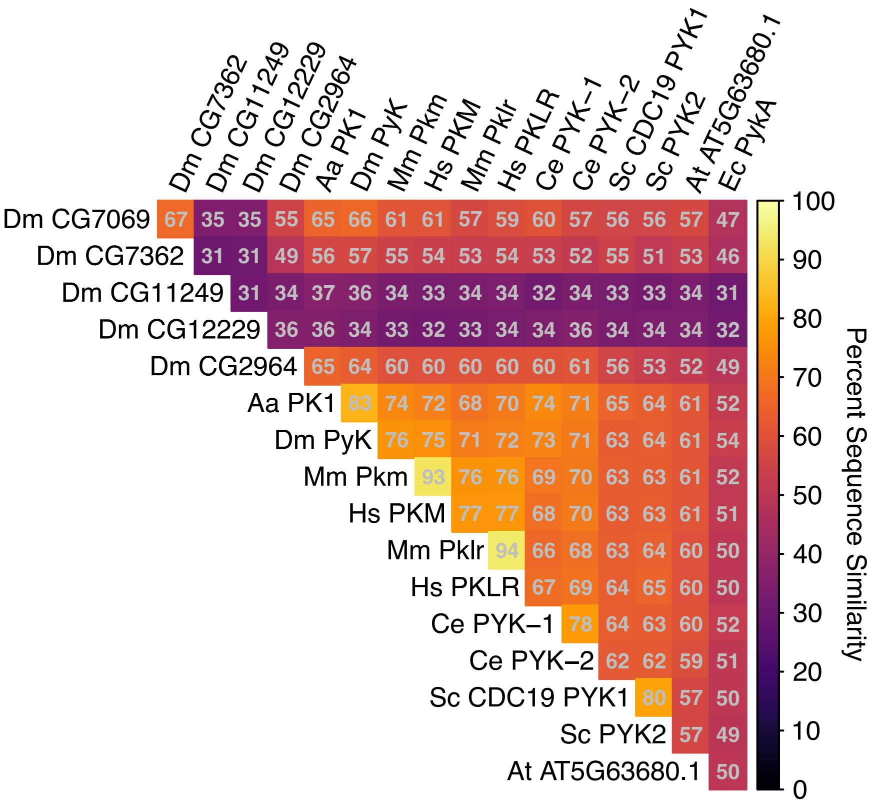


**Figure S2. A comparison of sequence similarities between Pyk homologs.** A heatmap of pairwise sequence identities extracted from the ensemble MSA. Species abbreviations: Aa, *A. aegypti; Dm*, *D. melanogaster*; Hs, *H. sapiens*; Mm, *M. musculus*; Ce, *C. elegans*; Sc, *S. cerevisiae*; At, *A. thaliana*; Ec, *E. coli.* See Supplemental Table 1 for a list of isoforms used in this analysis.
