## Supplementary material for "Metabolomic analysis of *Drosophila melanogaster* larvae lacking Pyruvate kinase": Figures S3

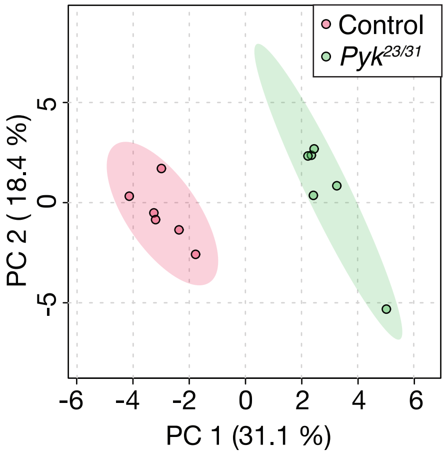


**Figure S3. A comparison of the metabolomic data from *Pyk*$^{23/31}$ mutant and *Pyk*$^{21/+}$ control samples using principal component (PC) analysis.** Targeted metabolomics data from Table S2 was analyzed using principal component analysis. Analysis was conducted using Metaboanalyst 5.0.
