## Supplementary material for "Metabolomic analysis of *Drosophila melanogaster* larvae lacking Pyruvate kinase": Figure S4

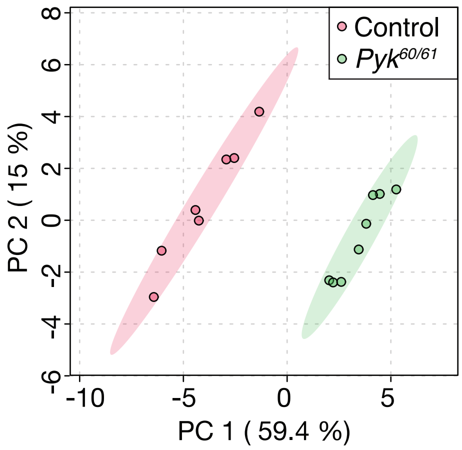


**Figure S4. A comparison of the metabolomic data from *Pyk*$^{60/61}$ mutant and *Pyk*$^{prec}$ control samples using principal component (PC) analysis.** Targeted metabolomics data from Table S3 was analyzed using principal component analysis. Analysis was conducted using Metaboanalyst 5.0.
