## Supplementary material for "Metabolomic analysis of *Drosophila melanogaster* larvae lacking Pyruvate kinase": Figure S5

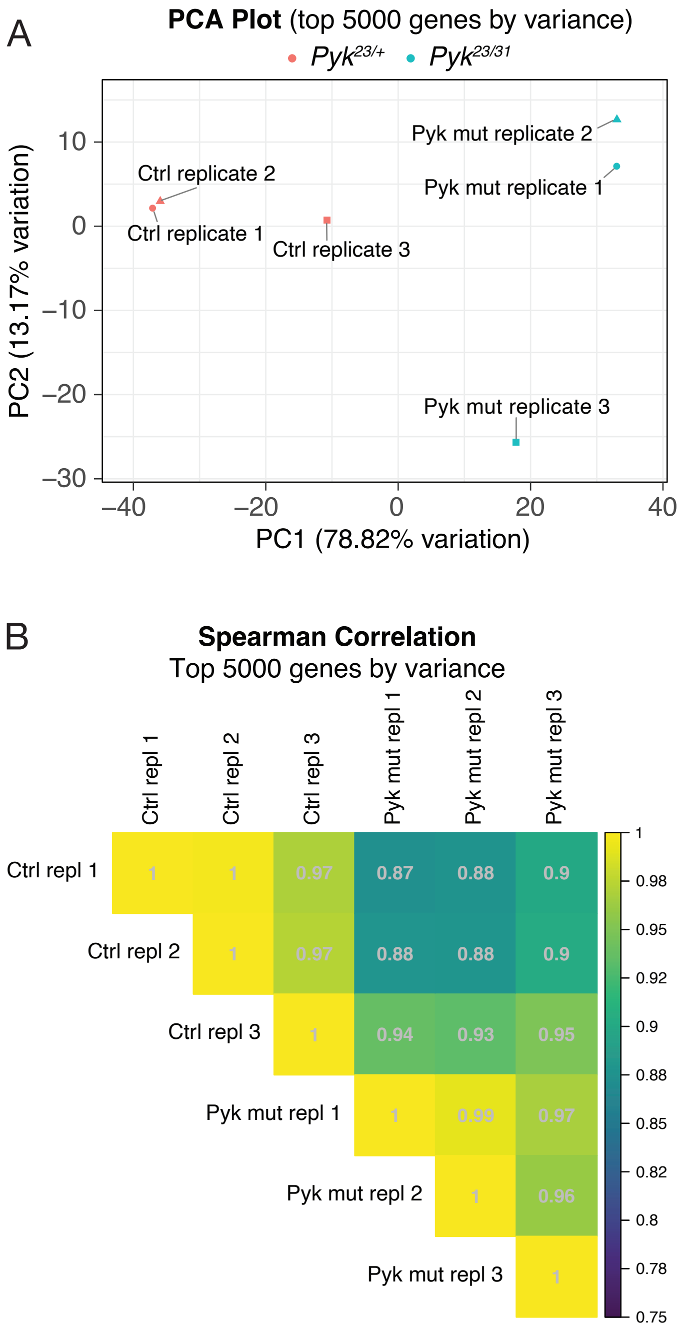


**Figure S5. Principal Component Analysis and Correlations of Experimental Replicates.** (A) PCA biplot of principal component 1 (PC1) versus PC2 for the top 5,000 genes filtered by variance with replicates labeled. (B) Spearman correlation plot of the top 5,000 genes filtered by variance after filtering out genes with either low expression levels or no detectable expression.
